# siRNA-based Therapeutic Candidate Targeting PRDM2 for Inhibition of Lung Cancer Progression

**DOI:** 10.1101/2025.02.06.636957

**Authors:** Sanjay Kumar, Md Zubbair Malik, Maya Chaturvedi, Mohit Mishra, Marzia Di Donato, Amelia Casamassimi, Ciro Abbondanza, Patrizia Gazzerro, Chuong Nguyen, Nicole N Nuñez, Somdutta Sen, Brad Niles, Rupesh Chaturvedi

## Abstract

Lung cancer is the leading cause of tumor-related fatalities worldwide. The current precision therapies are effective but target only those molecules that are impacted in a minority of lung cancer patients. Positive Regulatory Domain 2 (PRDM2), a member of the PRDM gene family, also known as Retinoblastoma Interacting Zinc finger (RIZ) protein. RIZ is found to be misregulated in more than 60% of lung cancer patients, representing a significantly affected population when compared to other mutation-prone oncogenic factors. RIZ expresses two contrasting variants, a tumor suppressor RIZ1, and an oncogenic RIZ2 protein. In lung cancer, the RIZ1/RIZ2 ratio alters, and the epigenetic silencing of RIZ1 promotes RIZ2 overexpression. In this study, we used siRNA (ARIZ-047) to knockdown and inhibit the effect of the oncogenic protein RIZ2. Inhibiting RIZ2 with ARIZ-047 treatment increased RIZ1 expression and decreased the viability of a lung cancer cell line (A549). The ARIZ-047 treatment also upregulated the negative regulators of the Wnt signaling pathway and restrained the tumor progression in the xenograft mice model. In conclusion, this study identifies ARIZ-047 as a targeted therapy in lung cancer, which specifically inhibits RIZ2 and suggests the role of the Wnt signaling pathway in RIZ2 overexpression.

**Graphical Abstract:** 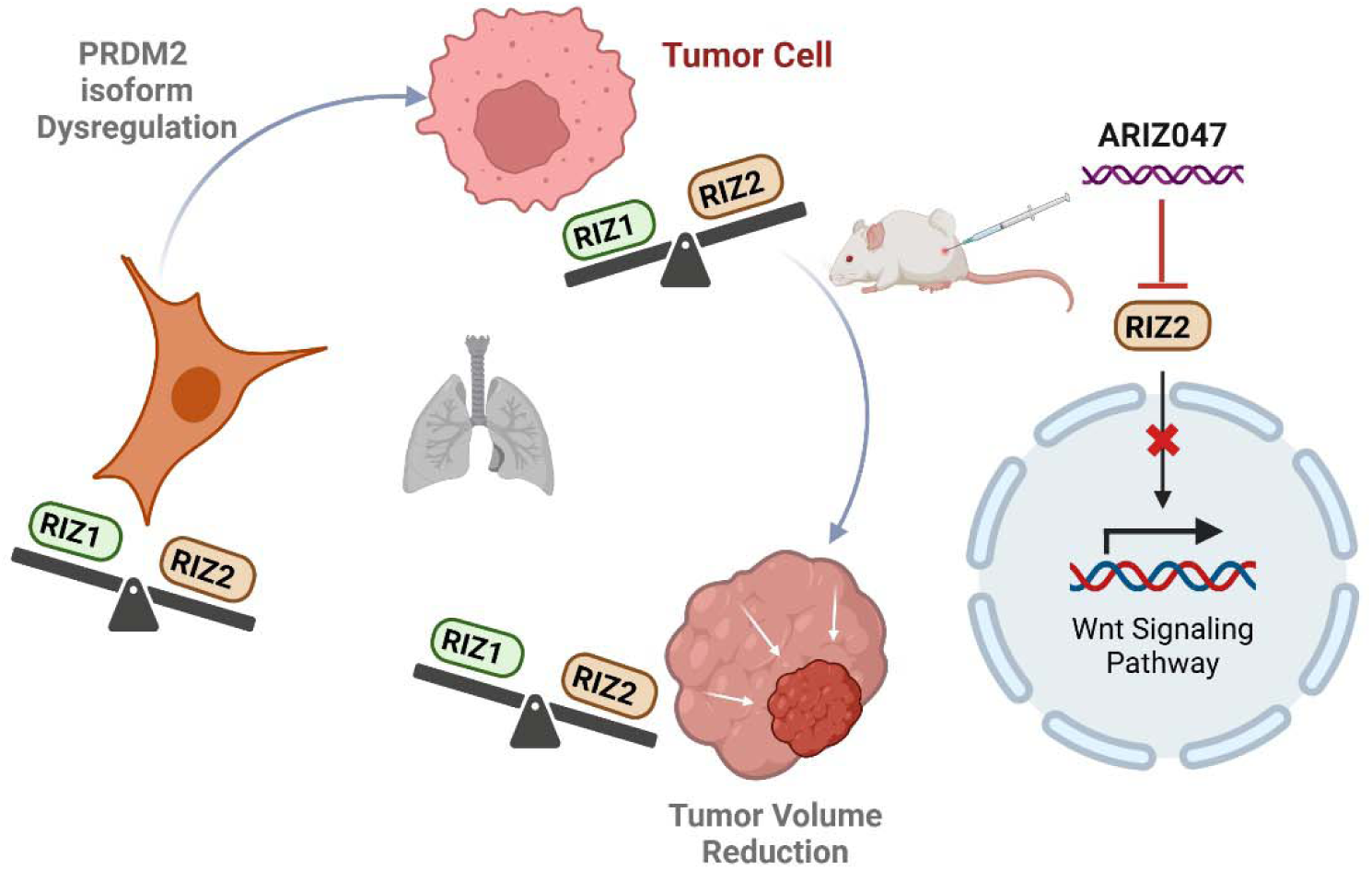

## Introduction

Lung cancer is the leading cause of cancer-related deaths worldwide, with an 18% mortality rate, followed by 13.6% and 9% for breast and colorectal cancer, respectively (1). Traditional cancer treatment modalities such as chemotherapy and radiation-based therapies have been pivotal in managing cancer; however, they are often associated with substantial side effects such as hematological and gastrointestinal complications along with impaired immune functions that can adversely impact patient’s quality of life (2,3). Although certain immuno-oncology approaches, including antibody therapy and chimeric antigen receptor (CAR-T) therapy, have exhibited efficacy in a limited range of patients, their widespread implementation is hindered by their substantial cost (4). Similarly, targeted therapies such as AstraZeneca’s Tagrisso have exhibited remarkable efficacy against EGFR-driven lung cancer but are confined to a small subset of patients with specific genotypes (5). Consequently, there exists a compelling rationale for exploring alternative therapeutic targets that encompass molecules affected in a large proportion of patients. The most common genetic mutations in lung cancers are reported to be approximately 50% in TP53, 30% in KRAS, and 5-10% in EGFR. (6), 5% in ALK (7), 3.5-4% in BRAF (8), and 1-4% in HER2 (9). Notably, PR/SET Domain 2 (PRDM2) emerges as a highly promising candidate, as it is disrupted in over 60% of lung cancer patients (10), thus positioning it as a potentially more effective therapeutic target.

PRDM2, also known as Retinoblastoma Interacting Zinc finger (RIZ) protein, is a prominent member of the PRDM gene family (11), which encompass a diverse group of transcription factors that collectively regulate critical cellular processes such as cell proliferation, differentiation, and fate determination (10). PRDM2 exists in two distinct protein forms, namely PRDM2a (RIZ1) and PRDM2b (RIZ2), distinguished by the presence or absence of the PR domain. Of particular significance, RIZ1, encompassing the PR domain, exhibits tumor-suppressive properties, highlighting its pivotal role in cancer regulation. Conversely, RIZ2 represents an oncogenic isoform lacking the PR domain, thus implicating its potential involvement in promoting tumorigenesis (10,12).

In normal tissues, the RIZ1 and RIZ2 express widely in similar ratios, and the malignancies could result due to the imbalanced expression of these two proteins (13). This imbalance favors the overexpression of the RIZ2 oncogenic protein, often caused by deletion, inactivating mutations, or silencing of the PR domain in the RIZ1 protein, leading to a decrease in its expression level. The overexpression of RIZ2 and the genetic or epigenetic silencing of RIZ1 have been observed frequently in human cancer cells and tissues (14). Genetic investigations on cancer cell lines demonstrated that genetic or epigenetic inactivation of the PR domain affects its histone methyltransferase activity, which plays a critical role in tumor suppressive action of RIZ1 (15). These studies suggest that RIZ1 opposes cell growth and tumorigenesis, whereas RIZ2, through its mitogenic properties, promotes cell proliferation and causes malignancies.

This study presents a comprehensive investigation of the therapeutic potential of ARIZ-047, an in-house developed siRNA-based treatment targeting the oncogenic protein RIZ2 in lung cancer. By employing a multi-faceted approach, we evaluated the effects of ARIZ-047. We found downregulation of RIZ2 and upregulation of tumor suppressor RIZ1 protein in the lung adenocarcinoma cell line (A549) without affecting the expression of off-target genes. Through the RNA seq experiment, we characterized the alterations in the global RNA landscape and identified regulatory changes in biological processes and signaling pathways. Additionally, we investigated the interplay between RIZ2 and Wnt signaling, specifically examining the influence of Wnt signaling on spheroid growth. Lastly, we evaluated the efficacy of ARIZ-047 in a xenograft mice model, assessing its significant potential for tumor size and volume reduction.

## Materials and Methods

### Cell Culture and Transfection

HEK-293 cells were cultured and transfected as described elsewhere (16). A549 cell line was procured from the National Center for Cell Science (NCCS), Pune, India. These cells were grown in Dulbecco’s Modified Eagle Medium (DMEM, Hi-media) enriched with 10% Fetal Bovine Serum (FBS, Hi-media), 2 mM L-glutamine (BD Biosciences), and 1% penicillin-streptomycin (Hi-media) under standard conditions at 37°C in a humidified condition containing 5% CO_2_. HBE1 cells were grown and maintained in Keratinocyte-Serum Free medium, enriched with 5 ng/ml human epidermal growth factor and 0.05 mg/ml bovine pituitary extract (Invitrogen, Cat. No. 17005-042). The media was also supplemented with 0.005 mg/ml of insulin and 500 ng/ml of hydrocortisone to ensure the optimal growth condition for HBE1 cells.

A549 and HBE1 cells were seeded at a density of 1 million cells per well in a 6-well plate for transfecting siRNAs and ScRNA using Lipofectamine reagent in Opti-MEM^TM^ Reduced Serum Medium (Thermo Fischer Scientific, Catalog number: 51985034) for 6 hrs. following the manufacturer’s instructions. For A549 cells, a Lipofectamine reagent from Dharmafect (Dharmacon) was used, whereas for HBE1 cells, a Lipofectamine reagent from RNAiMAX (Invitrogen) was used. 6 hrs., the Opti-MEM was replaced with a complete DMEM medium, and cells were allowed to grow for an additional 24 hours. Transfected cells were then ready for subsequent analysis.

### 3D Spheroid Culture

The 3D spheroids from HEK-RIZ2-GFP and HEK-GFP cells were prepared using the growth-factor reduced Matrigel Basement Membrane Matrix (BD Biosciences) described in our previous study (16). A549 cells were suspended in ice-cold Matrigel (Phenol red-free; Corning) and plated onto pre-warmed 96-well round bottom plates at a density of 2-32 cells per 50 µl of Matrigel per well. The Matrigel-cell mixture was allowed to polymerize for 15-20 minutes at 37°C. Subsequently, 200 µl of warm DMEM (Himedia) supplemented with 50 ng/ml human recombinant EGF (Sigma-Aldrich), 1 mM n-Acetylcysteine (Sigma), 1% N2 Supplement (Invitrogen), 2% B27 Supplement (Invitrogen), and 1% Pen/Strep (Invitrogen) was carefully added to each well. The plates were then incubated at 37°C with 5% CO2 for a period of 3 days to allow for cell growth and spheroid formation. ARIZ-047 treated cells were used for spheroid culture. Cells were embedded in Matrigel and cultured in a 3D culture medium with and without R-spondin (100ng/ml) for 6 days.

### In vitro cell viability assay

A549 and HBE1 cell viability was assessed by the 3-(4,5-dimethylthiazol-2-yl)-2,5-biphenyltetrazolium bromide (MTT) assay as previously described (17). The measurements were taken at 570 nm wavelength for absorbance and 690 nm for the background subtraction. The cell number was calculated as previously described (16).

### RNA extraction and qRT-PCR

Total RNA was extracted from the cells using Trizol solution (ThermoFisher Scientific), followed by the cDNA preparation using the previously described procedure (16). PGK1 was used as a housekeeping gene. Quantitative RT-PCR analysis was performed using the SYBR Green PCR Master Mix (New England Biolabs), 160 nM of each primer, and about 50ng of cDNA (RNA equivalent) as a template in an iCycler thermocycler (BioRad). PCR conditions were: Initial Denaturation at 95°C for 60 seconds, followed by 40 cycles of Denaturation at 95°C 15 seconds, then Extension at 60°C for 30 seconds. This was followed by performing a Melt Curve from 60-95°C. RIZ1 transcripts were amplified with forward (5’-CGTCTACCCTTCTCTGTGGA-3’) and reverse (5’-ATGAACCCTCAGTCTCAAGTG-3’) primers. RIZ2 transcripts were amplified with forward (CTGCTGACTTGAGTGAGAACA) and reverse (ATTTCGTATCCTTTGCCGTAGT) primers. KRAS transcripts were amplified with forward (5’-GCGTAGGCAAGAGTGCCTTG-3’) and reverse (5’-GTACTGGTCCCTCATTGCACTGTAC-3’) primers. ATP1A4 transcripts were amplified with forward (5’-CCACCCTAGGCTTCTCTCCT-3’) and reverse (5’-GCTATGGCCCTTTGTCAGGT-3’) primers. TOPBP1 transcripts were amplified with forward (5’-GGACAACCACTTCAGAAGGAGC-3’) and reverse (5’-CCTGGAGTTTCCAAGGCTGCAA-3’) primers. All reactions were carried out in triplicate for every cDNA template, and the melting curves were analyzed to verify the specificity of the reaction. The relative gene expression was calculated using the 2^ΔΔCt^ method (18). For normalization, the PGK1 gene was used.

#### Microscopy analysis

HEK293 cells stably expressing GFP and RIZ2-GFP were cultured in ultra-low attachment plates and allowed to grow in spheroid-promoting media. Spheroids were treated with sFRP3, WNT5, and LGK-974 for up to 12 days. Bright-field microscopy was used to capture images of spheroids at 4 days and 12 days post-treatment.

### RNA sequencing: Library Preparation, sequencing, and Quality control

According to the manufacturer’s protocol, the RNA-seq libraries were constructed using a NEBNext® Ultra™ RNA Library Prep kit (New England Biolabs, UK). RNA-seq library underwent paired-end sequencing (2 × 150 bp) using the Illumina HiSeq™ 2500 platform. The quality of the raw reads was assessed using FastQC and MultiQC. Adapter sequences were trimmed, and any low-quality reads or reads below a certain length from the raw FastQC reads were removed using Trimmomatic software. The clean reads were aligned to the hg19 assembly of the human reference genome using the Hisat2 alignment program. All sequence data were assessed for quality control using FastQC/MultiQC—quality control. The reference genomes used were Homo sapiens GRCh38 for human datasets. Accurate alignment was executed using HISAT2version 2.1.0. Assembly and quantification software packages, such as StringTie version 1.3.4d and Cufflinks version 2.2.1, were used. Samtoolsversion 1.9 was used for the file format conversion required during the alignment and quantification steps. The differentially expressed gene (DEG) analysis tools include Cuffdiff version 2.2.1., edgeR, and Limma.

### Gene Ontology & Pathway Enrichment Analysis

Gene ontology (GO) analysis was used to categorize genes into hierarchical categories and identify the gene regulatory network based on biological processes and KEGG pathways. DAVID bioinformatics tool (https://david.ncifcrf.gov/) was used to conduct gene ontology (GO) biological process and functional enrichment analysis of differentially expressed genes (DEGs). A p-value of <0.05 was recognized as significantly enriched. The statistically most enriched GO terms were visualized in ggplot2 (19).

### Construction of Protein-Protein Interaction (PPI) network

Using the GeneMANIA database, a PPI network for the DEGs was identified. A complex network in which proteins involved in a biochemical event interact is called the PPI network by considering the DEGs. The PPI network was constructed using Cytoscape v3.7.1 (20). Cytoscape software is a powerful tool for visualizing and analyzing PPI networks.

### PPI Network Involving Topological Analysis

The topological properties of the PPI network were analyzed using the Cytoscape plugins and Network Analyzer (21). The characteristics of the PPI network, including degree distribution (P(k)), clustering coefficient (c(k)), neighborhood connectivity (CN(k)), centrality betweenness (CB), and closeness (CC), were examined. The Degree (k), Betweenness centrality (C_B_), and Closeness centrality (C_C_) were measured as per the formulations described in our previous study (22).

#### Driver gene identification

The top ten genes with the highest degree, betweenness, and closeness centrality were used to extract driver genes. Common genes for the above-mentioned three topological properties were traced to the top ten hub genes (genes with maximum degrees) to ensure their reliability as driver genes of the regulatory network.

#### Weighted Co-Expression Network Analysis and Hub Module Identification

The Weighted Gene Co-expression Network Analysis (WGCNA) R package (23) was used to construct co-expression networks and classify module genes that are associated with clinical characteristics. DEGs and samples underwent quality filtering using the “good Samples Genes” function to handle missing values. Outliers were identified via sample clustering and removed. Statistical metrics such as mean expression per array and the count of missing values per array were documented. Low-variance genes and arrays with excessive missing data were excluded. The appropriate soft-thresholding power (β) was determined using the “pickSoftThreshold” function based on the scale-free topology criterion to enhance co-expression similarity in network construction. A weighted adjacency matrix was transformed into a topological overlap matrix (TOM), and dissimilarity (dissTOM) was computed. The “hclust” function was used to create a hierarchical clustering tree (dendrogram), and modules were identified using a dynamic tree-cut algorithm, which partitioned the dendrogram branches. Module Eigengene (ME) values were calculated, and modules with similar expression patterns were merged based on Pearson correlation. Absolute gene significance (GS) was computed to assess the correlation between modules and the trait of interest (weight). Module Membership (MM) was computed for all modules, representing the correlation between MEs and gene expression profiles. The correlation between MM and GS facilitated the identification of the most meaningful associations. Ultimately, the module displaying the highest and most significant correlation with weight was chosen as the central hub module.

#### Calcium Phosphate Nanoparticle loading

*Step 1: Aptamer PEGylation.* This step produces a DNA aptamer (GGTGGTGGTGGTTGTGGTGGTGGTGG) with PEG attached to the 5’ end of the DNA aptamer. Solutions of 0.431M methoxy-PEG-NH2 2000 (Laysan Bio) in 0.172M Imidazole HCL Sigma) in 1.26mL 1M MES (2-(N-Morpholino) ethanesulfonic acid: Sigma) buffer pH4, and 1mM DNA aptamer (e.g., AS1411) in water were prepared. In each tube, 50µl aptamer, 460µl nuclease-free water, 105µl MES/EDC, and 435µl PEG/Imidazole were mixed and incubated at 50°C under nitrogen gas overnight while stirring. The reaction mixture was slowly cooled and transferred to a 4 ml 3K MWCO filter pre-washed 4 times once with 0.05M NaOH, then 3 times with nuclease-free water at 7,500g. Nuclease-free water was added to the solution (1.5mL reaction mix with 2.5mL water). The aptamer solution was centrifuged at 7500g until the added water volume was in the flow through. This step was repeated four more times, increasing the water added and thus collected by 0.5mL each time (the last two cycles use the same volume). The mixture was collected and resuspended to the desired concentration. An EtBr agarose gel was run to determine the PEGylation percent by image densitometry.

*Step 2: siRNA PEGylation.* This step produces a siRNA with PEG attached to the 5’ end of the sense strand (5’P-GGGAGAGAfUGAGAGAGAAAdTdT). 0.431M methoxy-PEG-NH2 2000 (Laysan Bio) in 0.172M Imidazole HCL (Sigma) pH6, 1.26mL 1M MES (2-(N-Morpholino) ethanesulfonic acid: Sigma) buffer pH4, and 1mM of a siRNA sense strand were prepared. To make PEGylated sense strands, 50µl siRNA sense strand, 460µl nuclease-free water, 105µl MES/EDC, and 435µl PEG/Imidazole were mixed and incubated at 50°C under nitrogen gas overnight while stirring. To anneal the corresponding siRNA antisense strand (5’P-mUmUfUCfUCfUCUCAUC6C6CC**CdTdT** (bold=phosphorothioate)) to the pegylated sense strand, the reaction mixture with the siRNA sense strand was slowly cooled and 50µl 1mM of a corresponding siRNA antisense strand was added, solution was vortexed and incubated at 70°C for 20 minutes under nitrogen and stirring. The reaction mixture was slowly cooled over 2 hours. The reaction mixture was added to a 50mL tube, then 1mL of 5M sodium acetate and 13mL ethanol were added. The mixture was vortexed and incubated at −80C overnight. Tubes were vortexed and centrifuged at 15,000g at 4C for 90 minutes. The supernatant was removed and saved. 4mL −20C 75% ethanol was added to the tube and incubated for 90 minutes at −80C, then centrifuged at 15,000g at 4°C for 45 minutes. The supernatant was removed and saved; tubes were covered with a Kimwipe and left inverted at 4°C to dry overnight. The tubes were brought to room temperature, and samples were resuspended and vortexed with 100 µl nuclease-free water. An EtBr agarose gel was run to determine the PEGylation percent. Equal volumes of 1mM non-PEGylated sense and antisense RNA were mixed, heated at 70°C for 20 minutes, and slowly cooled. Annealed non-PEG siRNA was added to PEG-siRNA to achieve 12.5% PEGylation.

*Step 3: Calcium Phosphate nanoparticle assembly:* 25 mM Calcium Chloride (CaCl2) (Sigma) and 25 mM Sodium Phosphate (Na2HPO4) (Sigma) were made to a pH of 10. PEG-siRNA (12.5% PEGylated) and CaCl2 were mixed in solution to get a concentration of 200uM (12.5% PEGylated siRNA) and 5mM CaCl2. When included, PEG-aptamer (75% PEGylated) was added at 25uM. Na2HpO4 was diluted to get a 4.05mM concentration based on the total desired solution volume and was added last. Solutions were vortexed between additions. The solutions were prepared and incubated for 2 days at RT. After incubation, nanoparticle size was examined using Dynamic Light Scattering. A 500 µl 100K MWCO centrifugal filter was washed 4 times, once with 0.05M NaOH, then 3 times with nuclease-free water at 3,000g until most of the solution was passed through the filter. The nanoparticle solution was centrifuged with the pre-washed 100K MWCO filter at 3,000g until 90% of the solution volume had passed through the filter. The sample was collected and 25mM CaCl2 pH10 was added to increase the concentration of CaCl2 by 2mM, then vortexed. The solution was incubated at 37C for 2 hours, slowly cooled, vortexed, and incubated overnight at RT. The solution pH was measured, and nanoparticle size was examined by Dynamic Light Scattering. A syringe filter was pre-washed with 0.5ml water, and the sample was passed through the filter. Samples were collected for DLS and siRNA quantification.

#### Xenograft studies

The A549 Cell line was revived, passaged, and grown for 10 Days to achieve an adequate cell population prior to inoculation. Following their growth, Cells were mixed with Matrigel in a ratio of 1:1 to achieve a cell count of 7X10^6^ within 200 µl of volume. The cell-Matrigel mixture was injected subcutaneously into the dorsolateral right flank of 6–8-week-old NCr-Foxn1nu (NCR Nude Male) mice. Animals were monitored for tumor growth. Once the tumor volume reached around 100 mm^3^, animals were randomized based on tumor volume, ensuring the tumor volume remained constant. This categorization was done a day before treatment initiation, with each cage assigned its respective group label. In total, 48 mice were categorized into six groups, each containing eight. Group-1 served as the vehicle control; Group 2 received intravenous treatment of Calcium Phosphate (CaP) nanoparticle encapsulated ARIZ-047 (ARIZ-047/CaP); Group 3 received IV ARIZ-047/CaP treatment along with an intraperitoneal (IP) injection of carboplatin; Group-4 received IV treatment of Targeted Calcium Phosphate nanoparticle (T-CaP) encapsulated ARIZ-047 (ARIZ-047/T-CaP), where AS1411 aptamer was the targeting ligand; Group-5 received IV treatment of ARIZ-047/T-CaP along with the 30 mg/kg intraperitoneal (IP) dose of Carboplatin; Group-6 was administered with 30 mg/kg of Carboplatin. A 5% Dextrose solution dissolved in water was used as a vehicle system for ARIZ-047/CaP and ARIZ-047/T-CaP, whereas for Carboplatin, normal saline was used. The dose volume in both cases was kept fixed at 10 ml/kg. For treatment, both ARIZ-047/CaP and ARIZ-047/T-CaP Nanojacket were administered via IV route thrice a week for the initial two weeks and then reduced to twice a week for the next two weeks. Clinical observation of the animals was conducted daily, and tumor volume and body weight were recorded biweekly. On the third day of completion of the study, blood samples were collected to separate serum from each animal under the influence of mild isoflurane anesthesia using the retro-orbital technique. Animals were euthanized humanely via CO2 asphyxiation technique.

Post-euthanasia, the mice were dissected to retrieve vital organs such as the liver, lung, kidney, and tumor. Organ weights were noted, transferred to properly labeled Eppendorf tubes, and flash-frozen in liquid nitrogen. Serum was collected from whole blood by centrifuging at 4°C at 10,000 RPM for 10 minutes. After the experiment, all samples were stored at −80°C until further analysis.

#### Statistical analysis

All results are reported as mean ± Standard Deviation (SD). All experiments were conducted independently in triplicates, with three independent repetitions (n ≥ 9). Differences between the experimental groups were analyzed using the Student’s t-test. All statistical analysis was performed using GraphPad Prism. Finally, differences between the means were considered significant in decreasing order at \*\*\*\**p*<0.0001, \*\*\**p*=0.0005, \*\**p*≤*0.005, *p<0.05*.

## Results

### ARIZ-047 decreases RIZ2 expression level but increases RIZ1 expression in human cancer cell line

Gene Expression Profiling Interactive Analysis (GEPIA2) was used to investigate the expression of the PRDM2 transcript. This revealed significant downregulation of RIZ1 transcripts (PRDM2-001 and PRDM2-002) in tumor samples, suggesting its tumor-suppressive role. In contrast, RIZ2 transcripts (PRDM2-003 and PRDM2-004) remained unchanged between the tumor and normal samples (**Supplementary Figure 1**). Normally, RIZ1 and RIZ2 are expressed in a balanced ratio, essential for cell cycle regulation, and alterations such as RIZ2 overexpression may promote tumor formation (13). To delve deeper into the transcriptional alterations observed in cancer, we performed ratio analysis using the data from “The Cancer Genomics Atlas” (TCGA), a comprehensive resource that provides insights into the genomic changes that drive cancer. Surprisingly, our findings indicated that the abundance ratio of PRDM2-003 in the context of the total PRDM2 genes was higher in tumor samples. In contrast, the expression ratio of PRDM2-002 and PRDM2-003 remained similar in normal tissues (**Figure 1A**).

**Figure 1.**
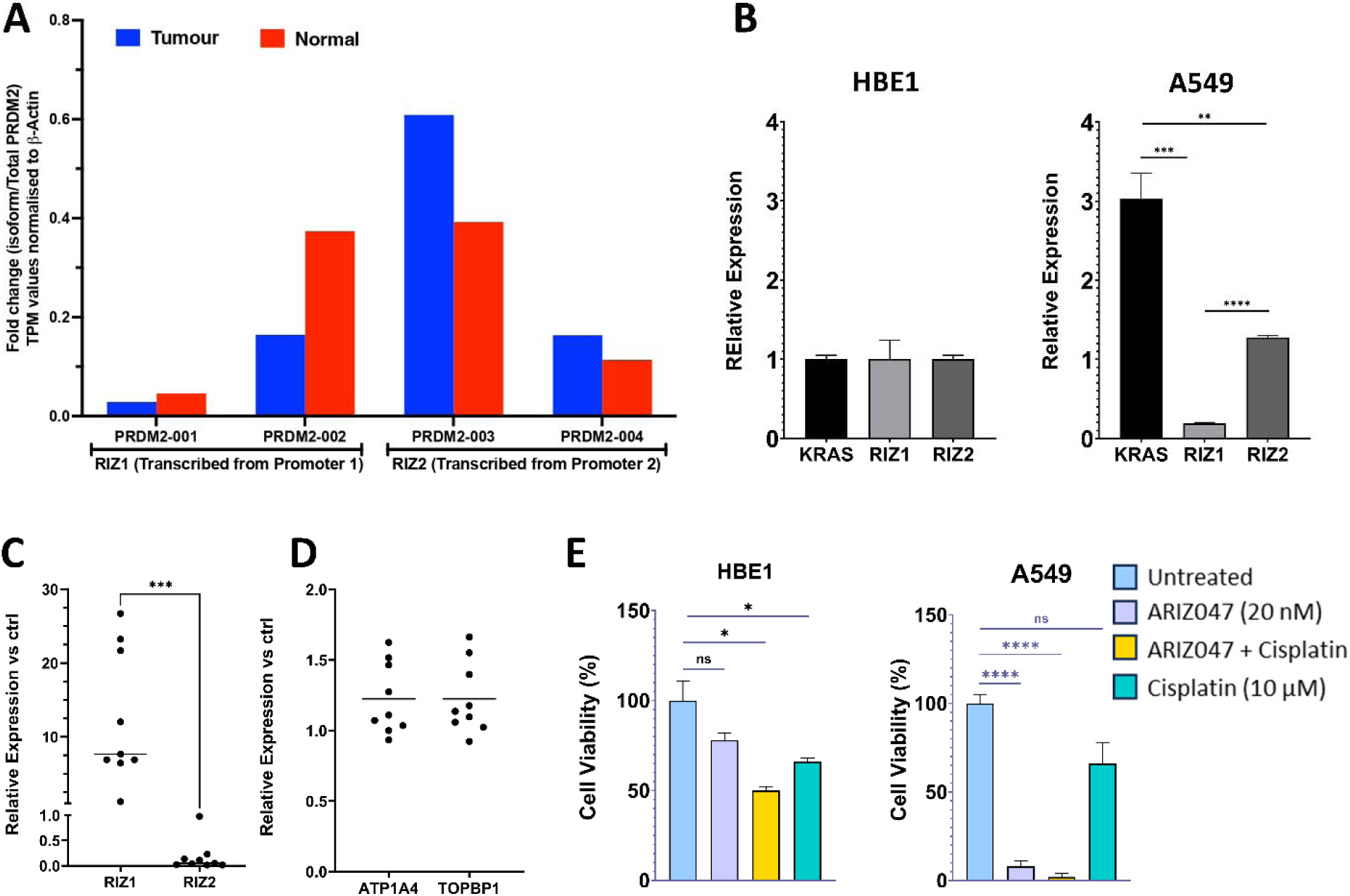
Effect of ARIZ-047 (20 nM) on RIZ1 and RIZ2 expression and cell viability. **A.** RIZ1 and RIZ2 expression data obtained from Cancer Genomics Atlas (TCGA), presented as fold change in TPM (Transcripts Per Million) values, normalized to β-actin. **B.** Expression of KRAS, RIZ1, and RIZ2 in HBE1 (non-cancer cell line) and A549 (lung cancer cell line). **C.** Relative expression of RIZ1 and RIZ2 in A549 cells treated with ARIZ-047 (RIZ2 siRNA) (***p<0.005). **D.** Relative expression of off-target genes, ATP1A4 and TOPBP1 in A549 cells treated with ARIZ-047. **E.** Cell viability analysis of cancer cell lines (A549) and non-cancer cell lines (HBE1) on treatment with ARIZ-047, ARIZ-047 + Cisplatin, and Cisplatin compared to untreated cells (*p≤0.01). Data represent ± SEM. *,p≤0.01; **,p=0.006; ***,p=0.001; and ****p<0.005.

The relative expression of tumor suppressor RIZ1 and oncogenic RIZ2 gene in a lung cancer adenocarcinoma cell line (A549) were compared to those in a non-cancerous Human Bronchial Epithelial (HBE1) cell line using quantitative real-time (qRT-PCR). The qRT-PCR analysis showed downregulation of RIZ1 and upregulation of RIZ2 in A549 cells, whereas HBE1 cells expressed a similar level of RIZ1 and RIZ2. Moreover, KRAS, a frequently mutated oncogene in non-small cell lung cancer (NSCLC), constitutes approximately 30% of all lung malignancies (24) was upregulated in A549, which was relatively higher than RIZ1 and RIZ2 (**Figure 1B**). These findings suggest that dysregulation of RIZ1 and RIZ2 may play a role in the development and progression of lung adenocarcinoma. Treatment with ARIZ-047 led to a significant reduction in RIZ2 and an increase in RIZ1 expression in A549 cells **(Figure C)**, demonstrating high specificity for RIZ2 with no significant changes in off-target genes ATP1A4 and TOPBP1 **(Figure 1D)**. When evaluating the specificity of ARIZ-047, HBE1 cells treated with the same ARIZ-047 concentration showed minimal cytotoxicity, with no significant reduction in viability, while cisplatin reduced viability in HBE1 cells by approximately 30%. Additionally, ARIZ-047 reduced A549 cell viability and proliferation by over 60%, outperforming cisplatin, which caused about a 40% reduction. These findings suggest that ARIZ-047 may have a therapeutic advantage over traditional chemotherapy by selectively targeting cancer cells while sparing normal cells (**Figure 1E)**. These results support the potential utility of ARIZ-047 as a targeted therapeutic agent for lung cancer, warranting further investigation into its clinical application.

### ARIZ-047 changes the RNA landscape and inhibits the Wnt Signaling pathway

To investigate the effect of ARIZ-047 on A549 cells, we performed a DEG analysis between ARIZ-047 and ScRNA-treated cells. Raw expression data processing resulted in 40500 unique genes after consolidating duplicate gene expressions (**Figure 2A, B**). A stringent filtering process excluded genes with low counts per million, retaining 1097 mRNAs followed by Trimmed Mean of M-value normalization and log transformation using the edgeR package. The Limma package identified 111 DEG (p < 0.05, absolute log2 fold change > 0.5), with 61 downregulated and 50 upregulated genes. These findings suggest that ARIZ-047 induces significant transcriptional changes in A549 cells, providing insights into its therapeutic effects in lung cancer. GO term enrichment analysis identified key biological pathways associated with **DEGs** in ARIZ-047-treated cells (**Figure 2C-D).** The downregulated DEGs were overrepresented in cell migration, JAK-STAT, cell proliferation, and antiapoptotic pathways, all implicated in tumor progression and metastasis (25). Conversely, upregulated DEGs were enriched in negative regulation of Wnt signaling and cell proliferation, along with positive regulation of p53 and type-I interferon signaling, which are critical in tumor suppression (26).

**Figure 2.**
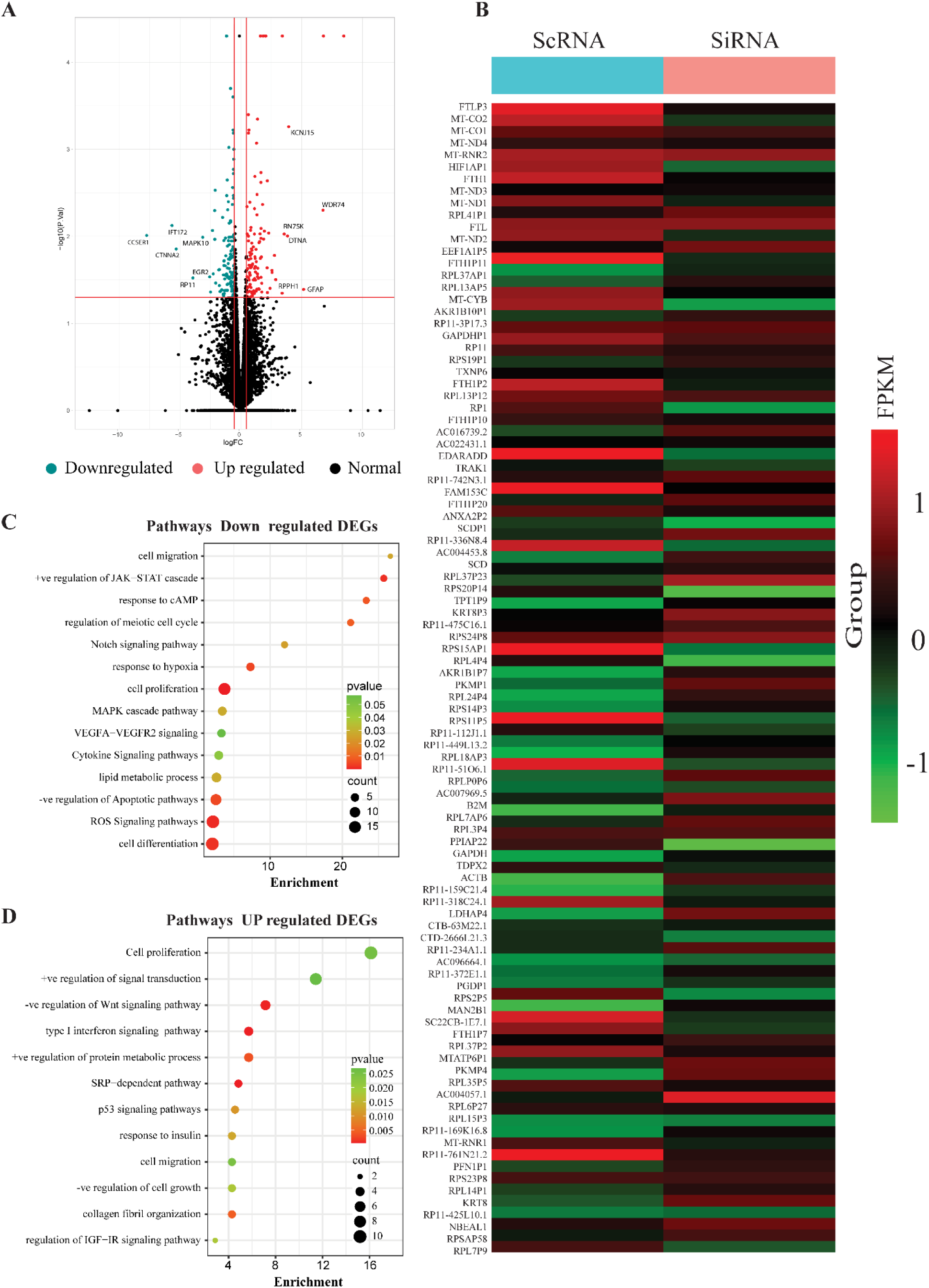
**A.** Volcano plot showing the variation of significance and log-fold-change for ARIZ-047 mediated’ differential gene expression analysis. **B.** The top 100 highly significant expression profiles are shown in the heatmap. **C.** Gene ontology (GO) enrichment analysis of ARIZ-047 mediated downregulated genes in anti-cancer relevant biological process. **D.** Gene ontology (GO) enrichment analysis of ARIZ-047 mediated up-regulated genes in anti-cancer relevant biological process.

### In silico addition of RIZ2 rearranges the protein-protein interaction network and recruits tumorigenic proteins as top hub proteins

To elucidate the intricate molecular mechanisms and cellular pathways underlying the progression of lung cancer in the presence and absence of the RIZ2 oncogenic protein, we conducted a comprehensive PPI network analysis. The ARIZ-047-induced changes in the RNA landscape showed global connectivity patterns of DEGs in the PPI network for various biological processes. Based on the degree, betweenness, and closeness centrality measures, top hub DEGs showed significant interactions with each other when the PPI network was formed (**Figure 3A**). A protein with a high degree (k) interacts with many other proteins in the PPI network (**Figure 3B**). Betweenness centrality (C_B_) (**Figure 3C**) and closeness centrality (CC) (**Figure 3D**) are topological metrics that demonstrate how efficiently signals move in a PPI network. The top hub DEGs identified in the PPI network, such as JUN, FOS, CYR61, TRIB1, KLF10, EGR1, KLF6, ATF3, BTG2, and IER2, are related to cellular proliferation, apoptosis, metastasis, and tumor development.

**Figure 3:**
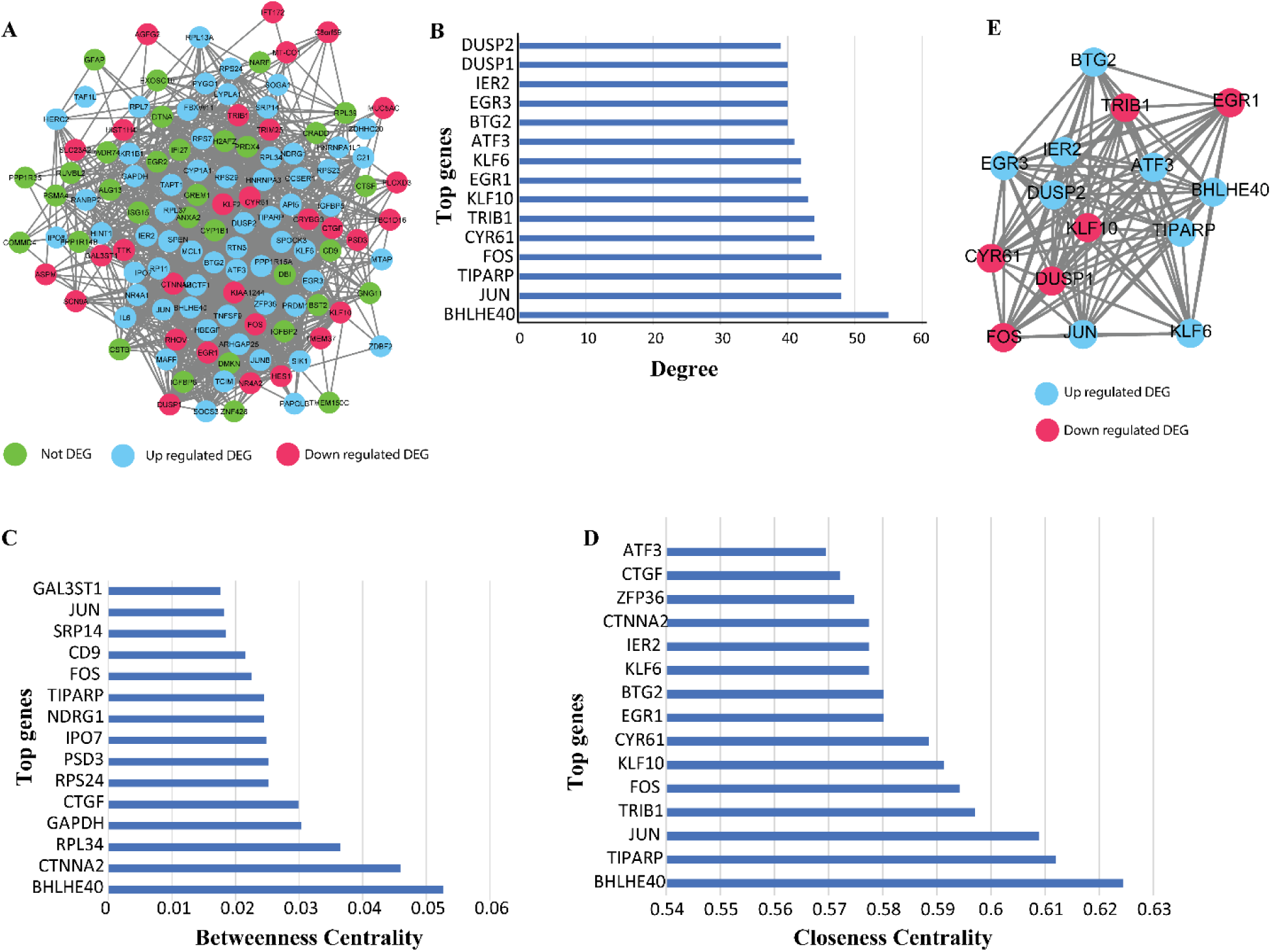
Protein-protein interaction (PPI) network prepared in the absence of RIZ2. **A.** Illustration of a PPI -Network showing Up-regulated and Down-regulated genes in the absence of RIZ2. **B.** The degree centrality, **C.** The betweenness centrality, and **D.** The closeness centrality of top hub proteins in the RIZ2 lacking PPI-network. **E.** The PPI network was prepared using the top hub proteins, which lacked RIZ2.

Furthermore, we observed significant global modifications in the interacting protein networks by adding RIZ2 protein in the PPI network (**Figure 4A**). Several top hub proteins selected based on the degree, betweenness, and closeness centrality exhibited altered interactions and connectivity patterns (**Figure 4B-D**). By *in silico* addition of RIZ2 in the PPI network, several top-hub proteins such as PRDM1, ZFP36, DUSP1, and MCL1 formed direct interaction with the RIZ2 protein (**Figure 4E**).

**Figure 4:**
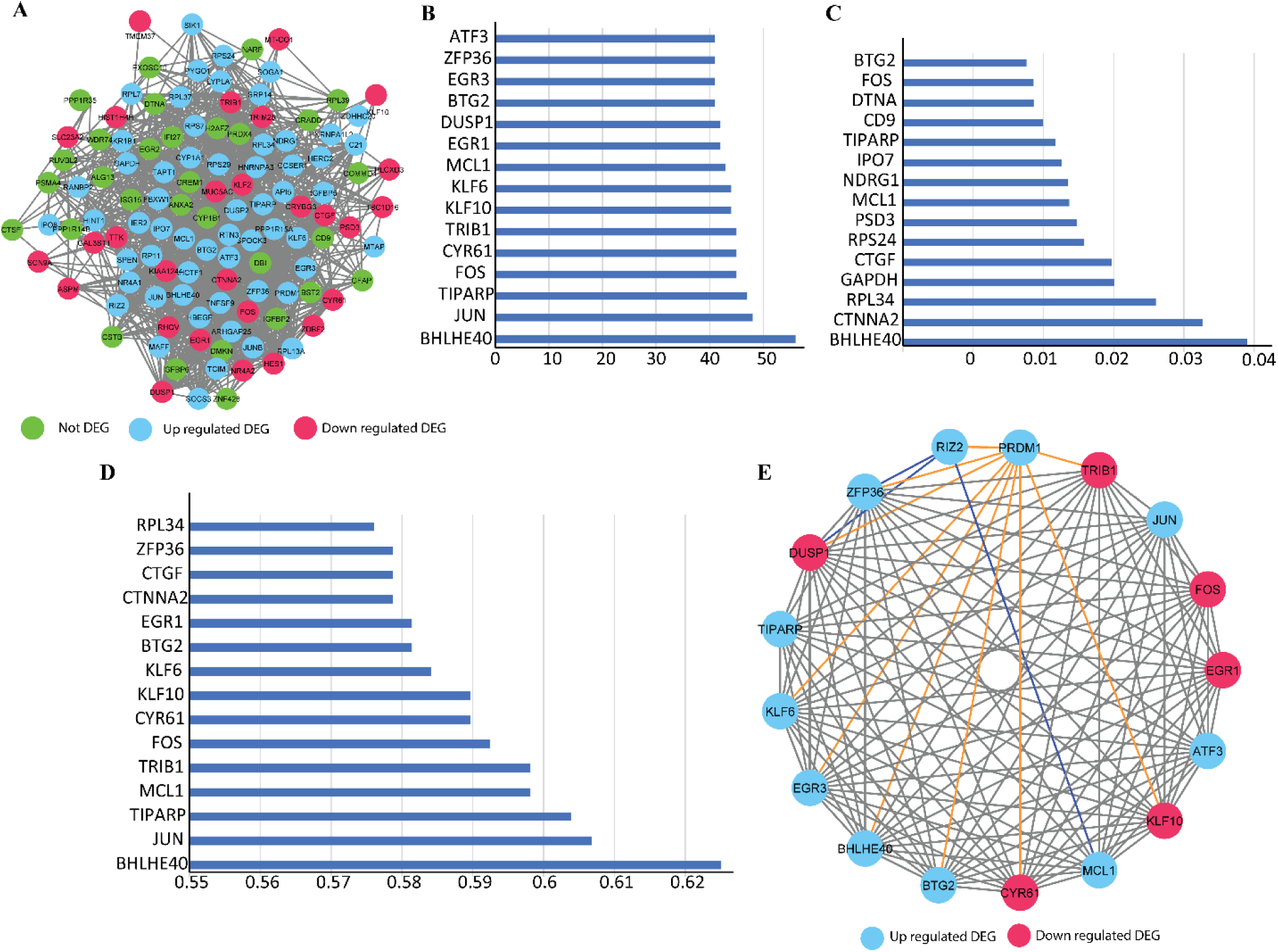
Protein-protein interaction (PPI) network prepared in the presence of RIZ2. **A.** Illustration of a PPI - Network showing Up-regulated and Down-regulated genes in the presence of RIZ2. **B.** Top degree centrality, **C.** The betweenness centrality, and **D.** The closeness centrality of proteins in the PPI network possessing RIZ2. **E.** Regulatory network of top hub proteins prepared in the presence of RIZ2.

Further, the PPI network consisting of RIZ2 was segregated into four distinct communities, each representing specific functional themes. The first community was enriched in DEGs involved in biological processes such as RNA binding, protease binding, Lamin binding, and integrin binding (**Supplementary Figure 2A**). The second community consisted of DEGs associated with translation and transcription (**Supplementary Figure 2B**). The third community comprised DEGs involved in cancer-associated signaling pathways such as cell proliferation, Wnt signaling, and apoptotic pathways (**Supplementary Figure 2C**). Lastly, the fourth community encompassed DEGs associated with metabolic pathways and immune responses (**Supplementary Figure 2D**).

### In silico deletion of top hub proteins negatively affects the connectivity and signaling within the protein-protein interaction network

The validation was essential for checking the healthiness of the obtained PPI network, and therefore, centrality values for the degree (k), betweenness (BC), and closeness (CC) in the context of RIZ2 absence and presence were determined (**Table 1, and Supplementary Figure 3**). Degree refers to the number of interactions a protein makes with other proteins in the PPI network. The Degree of the PPI network improved when RIZ2 was added to the ARIZ-047-treated PPI network. The BC value of 0.35 for the ARIZ-047 treated PPI network suggests a moderate signaling or information flow level within the network. The ARIZ-047 treated network, in addition to RIZ2, showed a BC value of 0.38, which suggests that the PPI network has a slightly higher level of signaling where RIZ2 might be serving as a more prominent bridge or mediator, facilitating information flow and communication between different proteins in the network. The CC value of 0.16 for the ARIZ-047 treated PPI network suggests that the average distance between two proteins in the PPI network is longer. In contrast, the ARIZ-047 treated PPI network added with RIZ2 protein showed a CC value of 0.3, which indicates two proteins are positioned closer, resulting in a shorter average distance. This implies that the ARIZ-047-treated PPI network containing RIZ2 protein signals can be transmitted more efficiently. Deletion of the top 15 hub proteins and then the top 50 hub proteins from both the ARIZ-047 responsive PPI network and the ARIZ-047 responsive PPI network added with RIZ2 resulted in the subsequent decrease in the value of degree, BC, and CC. It suggests these proteins are crucial in maintaining connectivity, information flow, and efficient communication within the PPI network. Their removal disrupts the network structure and potentially impacts the overall functionality and dynamics of the network.

**Table 1.** Topological parameters of protein-protein interaction network.

|  | Topological parameters of ARIZ-047 treated PPI network |  |  | Topological parameters of RIZ2+ARIZ-047 treated PPI network |  |  |
| --- | --- | --- | --- | --- | --- | --- |
|  | Degree | Betweenness | Closeness | Degree | Betweenness | Closeness |
| With all Top hub proteins | 0.1 | 0.35 | 0.16 | 0.47 | 0.38 | 0.3 |
| Deletion of top 15 Top hub proteins | 0.05 | 0.2 | 0.09 | 0.04 | 0.36 | 0.23 |
| Deletion of top 50 Top hub proteins | 0.023 | 0.08 | 0.03 | 0.01 | 0.21 | 0.1 |

### Identification of driver genes via weighted co-expression network analysis

The expression data encompassing 630 DEGs and their respective attributes were employed as input for the WGCNA analysis. Following the procedure detailed in the materials and methods section to determine the soft threshold (β) using WGCNA, an intriguing observation emerged, depicted in **Figure 5**. Figures 5A and 5B notably demonstrate a transition occurring at β=140 with an associated R2 value of 0.45. Considering this, we opted to utilize a power value of β=150 to construct the adjacency matrix for the weighted network. Subsequently, the biologically meaningful measure of node similarity, known as TOM, was computed from the adjacency matrix. Leveraging TOM as the distance metric, a hierarchical clustering of genes was performed, and the resulting dendrogram led to the identification of gene modules. The gene modules were delineated by setting a height cut-off of 0.90, guided by insights from the TOM plot (**Figure 5C**). This step was reiterated, yielding nine significant modules, each marked by a distinct color code, showcasing genes that are co-expressed in a coordinated manner. The outcome of the hierarchical clustering tree, facilitated by the dynamic tree cut algorithm, unveiled a comprehensive collection of nine color-coded gene modules (Black, Blue, Brown, Green, Pink, Red, Tortoise, and Yellow), as visually illustrated in **Figure 5C**. The modules’ sizes are as follows: black=49, blue=77, brown=72, green=63, green, yellow=74, pink=45, red=52, turquoise=92, and yellow=69. Each module represents various biological pathways that are involved in carcinogenesis. Our WGCNA analysis showed a community of different driver genes, of which 18 (nine upregulated and nine downregulated genes) were found to be differentially regulated with ARIZ-047 (**Figure 5D**). The centrality parameter analysis revealed five driver genes (**Figure 5E**). Based on the WGCNA and centrality parameter analysis, the EGR1 gene was the common driver gene. Furthermore, the interaction of EGR1 was analyzed with all PPI communities, where it was found that EGR1 made a maximum node connection with module 3, which includes many genes involved in cancer and Wnt-signaling pathways.

**Figure 5.**
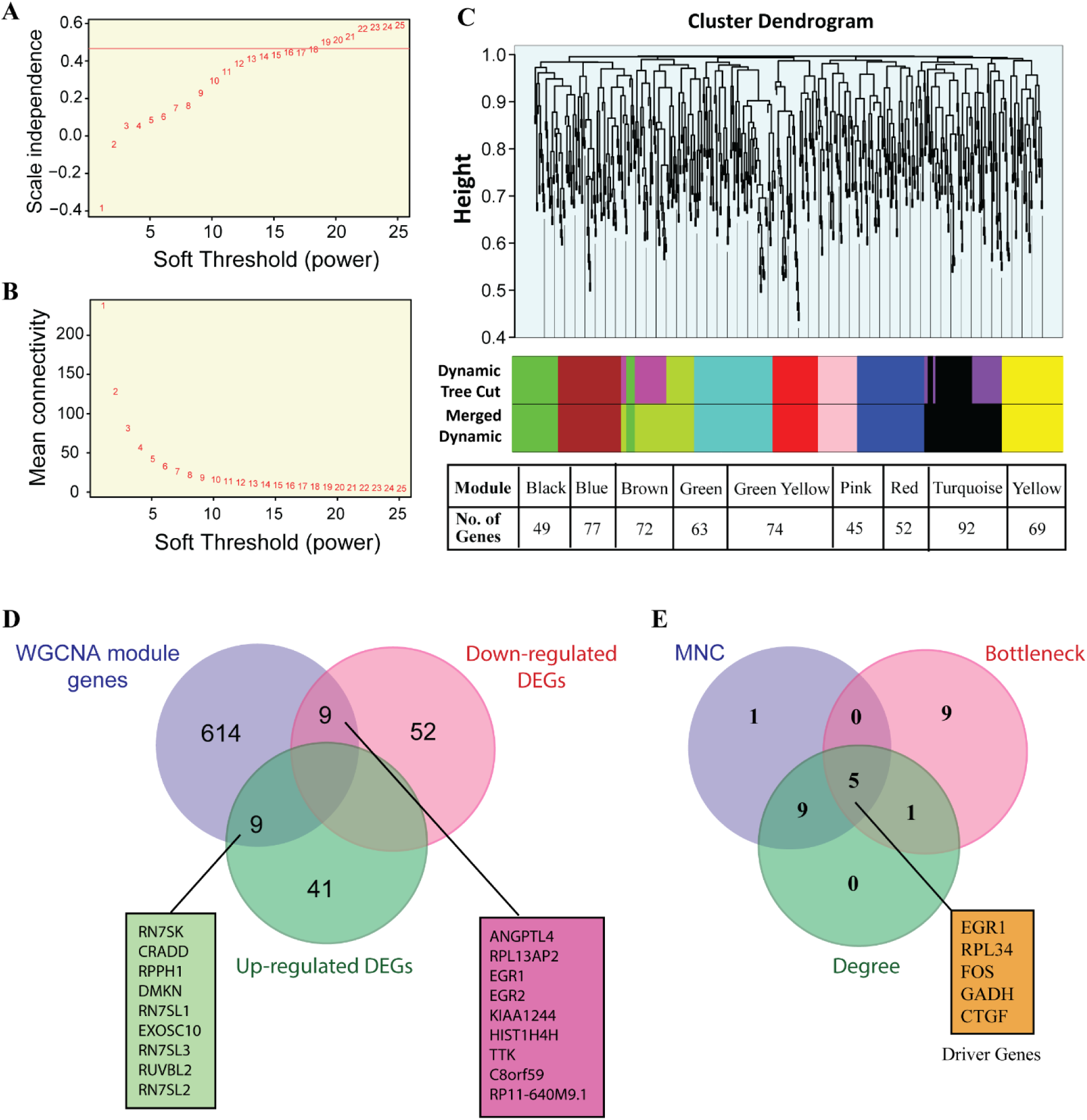
Weighted correlation co-expression network analysis of gene expression RNA seq dataset **(A)** Analysis of the scale-free fit index for various soft-thresholding powers. **(B)** Analysis of the mean connectivity for various soft-thresholding powers. **(C)** Gene dendrogram was obtained by clustering the gene expression from the dataset. Nine modules (Black, Blue, Brown, Green, Pink, Red, Tortoise, and yellow) were marked with different colors. **(D)** Common driver genes regulate WGCNA, down-regulated DEGs, and up-regulated DEGs. **(E)** Common driver genes are identified based on the centrality parameter analysis.

### RIZ2 modulates the WNT signaling pathway

To recapitulate the intricate in vivo architecture, 3D spheroids were successfully established from A549 and HEK293 cells. To investigate the impact of ARIZ-047 on cell viability and survival, we treated A549 Matrigel 3D lung spheroids with scRNA (control), ARIZ-047 (RIZ2 siRNA), and a combination of ARIZ-047 and R-spondin (Wnt signaling agonist). Microscopic images obtained over 6 days revealed that A549 cells treated with ARIZ-047 failed to form 3D organoids compared to those treated with control ScRNA. Notably, co-treatment with the Wnt signaling agonist R-spondin reversed the inhibition of 3D spheroid formation of A549 cells (**Figure 6A-B**), suggesting that ARIZ-047 inhibits the Wnt signaling pathway, which might be associated with RIZ2 expression.

**Figure 6.**
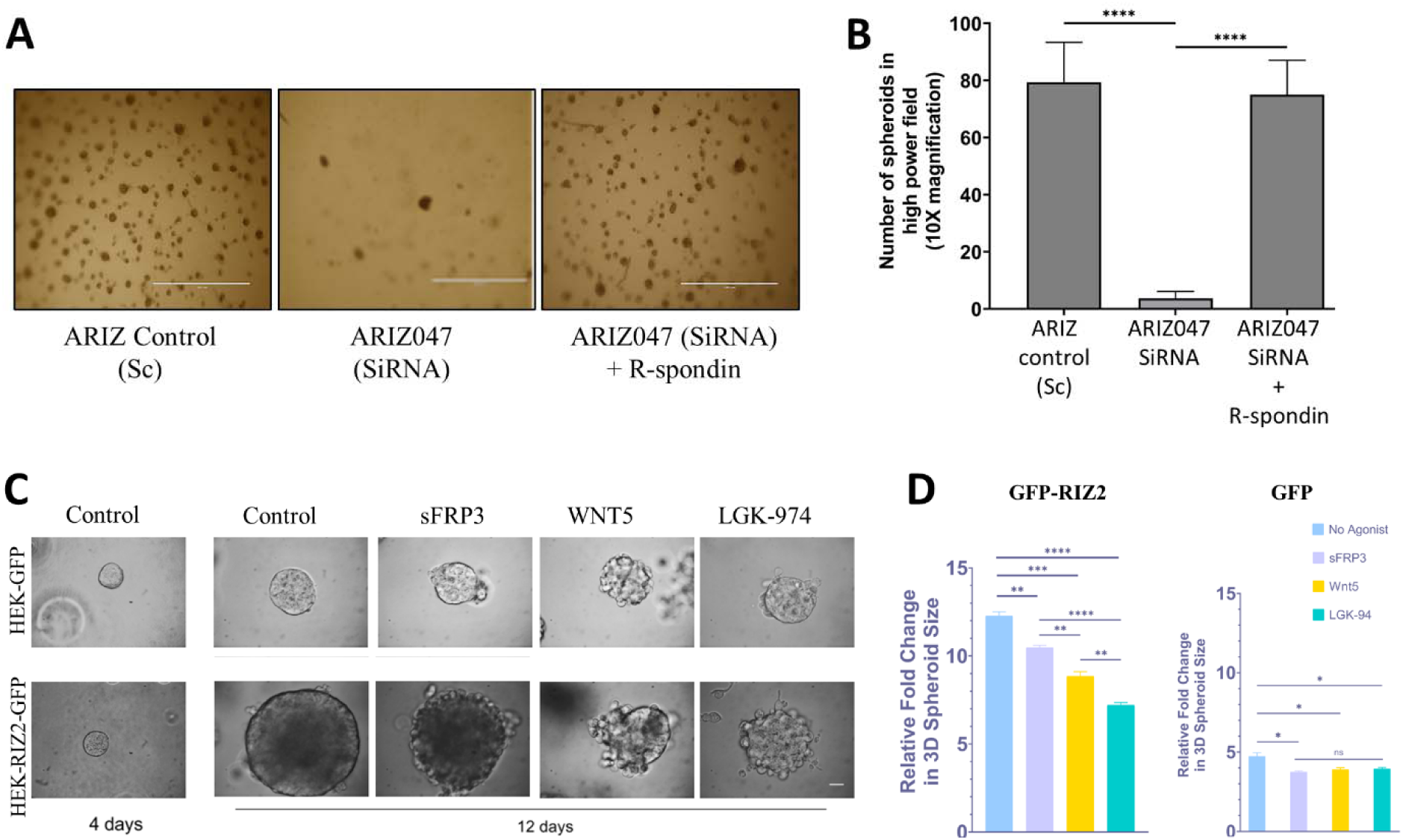
Potent inhibition of A549 Matrigel 3D lung cancer spheroids by ARIZ-047 is dependent on inhibition of WNT signaling. **A.** A549 cells were treated with ScRNA (ARIZ control), ARIZ-047 (RIZ2 SiRNA), and ARIZ-047 + R-Spondin, and their microscopic images were obtained after 6 days. **B.** Comparison of the number of spheroids formation on treatment with ScRNA, ARIZ-047, and ARIZ-047 + R-Spondin in high power field (****p < 0.0001). **C.** HEK-293 cells stable transfected with GFP and RIZ2-GFP were used to see the effect of Wnt-signaling agonist of the growth of miniatured 3D spheroids in ECM. **D.** Bar graphs representing the relative fold change in RIZ2 lacking- and overexpressing-spheroid size (area) in the presence of Wnt-signaling agonists, which was calculated using the Application Suite Software (****p<0.0001, ***p=0.0005, **p≤*0.005, *p<0.05*).

We further decided to investigate the effect of Wnt signaling pathway agonists such as sFRP3 (27), WNT5 (28), and LGK-974 (29), on 3D spheroids overexpressing the RIZ2 protein derived from HEK cells (lower panel) compared to control spheroids RIZ2 (upper panel), as shown in **Figure 6c**. Notably, after four days, phase contrast microscopic images obtained at 4 days showed the formation of well-defined roundish and well-differentiated 3D spheroids in both GFP and RIZ2-GFP overexpressing HEK-293 cells cultured in Matrigel. Subsequently, changes in the size and morphology of the spheroids overexpressing RIZ2-GFP and only GFP were monitored over an additional 12-day period in the presence of Wnt signaling agonists. Interestingly, an increase of about 12-fold in the size of pEGFP RIZ2-derived organoids was observed after 12 days, accordingly with the previous study (30).

After 12 days of culturing, spheroids displayed a moderate increase in size, whereas spheroids expressing RIZ2 exhibited a significant increase in size. Intriguingly, when RIZ2-overexpressing spheroids were subjected to the treatment with a Wnt signaling agonist, a substantial size reduction was observed compared to the untreated spheroids. Among all the Wnt-signaling agonists, LGK-974 significantly reduced the size of the RIZ2-GFP overexpressing spheroids. In contrast, the spheroids derived from GFP-overexpressing HEK-293 did not exhibit any alterations in size when exposed to sFRP3, WNT5, and LGK-974 (**Figure 6C-D**). These findings suggest RIZ2 might play a pivotal role in modulating WNT signaling and contribute to Lung cancer tumor progression. Further investigations are warranted to elucidate the precise molecular mechanisms underlying the interplay between RIZ2 and WNT signaling in the context of lung cancer.

### ARIZ-047 inhibits tumor growth in vivo

To assess the therapeutic potential of ARIZ-047 in vivo, we employed the NCR-Foxn1nu mice xenograft tumor model. Mice were divided into six groups, with eight mice per group based on their treatment (**Figure 7**). Mice bearing tumors were treated with various combinations with and without ARIZ-047, and changes in RIZ2 expression were measured by fold change compared to the control group. In the xenograft tumor model, administering ARIZ-047/T-CaP via intravenous (IV) route (Group 4) resulted in a significant two-fold reduction in RIZ2 expression compared to the vehicle disease control group (Group 1). RIZ2 expression was also observed in Groups 2, 3, 5, and 6, but the reduction was not as pronounced as in Group 4. Notably, Group 6, which received carboplatin, an FDA-approved anti-cancer drug, did not show a significant difference in RIZ2 expression compared to Group 4 (**Figure 7A**). These findings highlight the potent effect of the ARIZ-047/T-CaP combination in suppressing RIZ2 expression in the xenograft tumor model, surpassing the impact observed with carboplatin treatment alone. Tumor weight was assessed in all experimental groups, revealing that Group 4 exhibited the lowest average tumor weight at 234.21 mg. In contrast, the control group displayed a significantly higher tumor weight of 547.68 mg, while the remaining groups exhibited tumor weights ranging from 300 to 450 mg (**Figure 7B**). These results indicate that treatment in Group 4 substantially reduced tumor size compared to the control and other treatment groups. The ARIZ-047 treatment strategy is demonstrated in **Figure 7C**. The group treated with a dose of 1 mg/kg of ARIZ-047 exhibited a remarkable reduction in tumor volume compared to the control group and the group treated with 30 mg/kg carboplatin (**Figure 7D**). Notably, even at a substantially lower dose of 1 mg/kg, ARIZ-047 demonstrated significant efficacy in reducing tumor volume, surpassing the effectiveness of the 30 mg/kg carboplatin treatment. The ARIZ-047 was administered IV two to three times a week, whereas carboplatin was administered IP two times a week. These findings highlight the potent anti-tumor activity of ARIZ-047 and its potential as a superior therapeutic agent compared to carboplatin at higher doses. **Figure 7E** presents the representative images of mice from Group 1 (untreated) and Group 4 (ARIZ-047/T-CaP -treated), along with their corresponding extracted tumors. It is evident from the images that the untreated Group 1 mice exhibit larger tumor sizes compared to the Group 4 mice, which underwent treatment with ARIZ-047/T-CaP. These visual observations further support the effectiveness of ARIZ-047 in reducing tumor growth in the experimental model.

**Figure 7.**
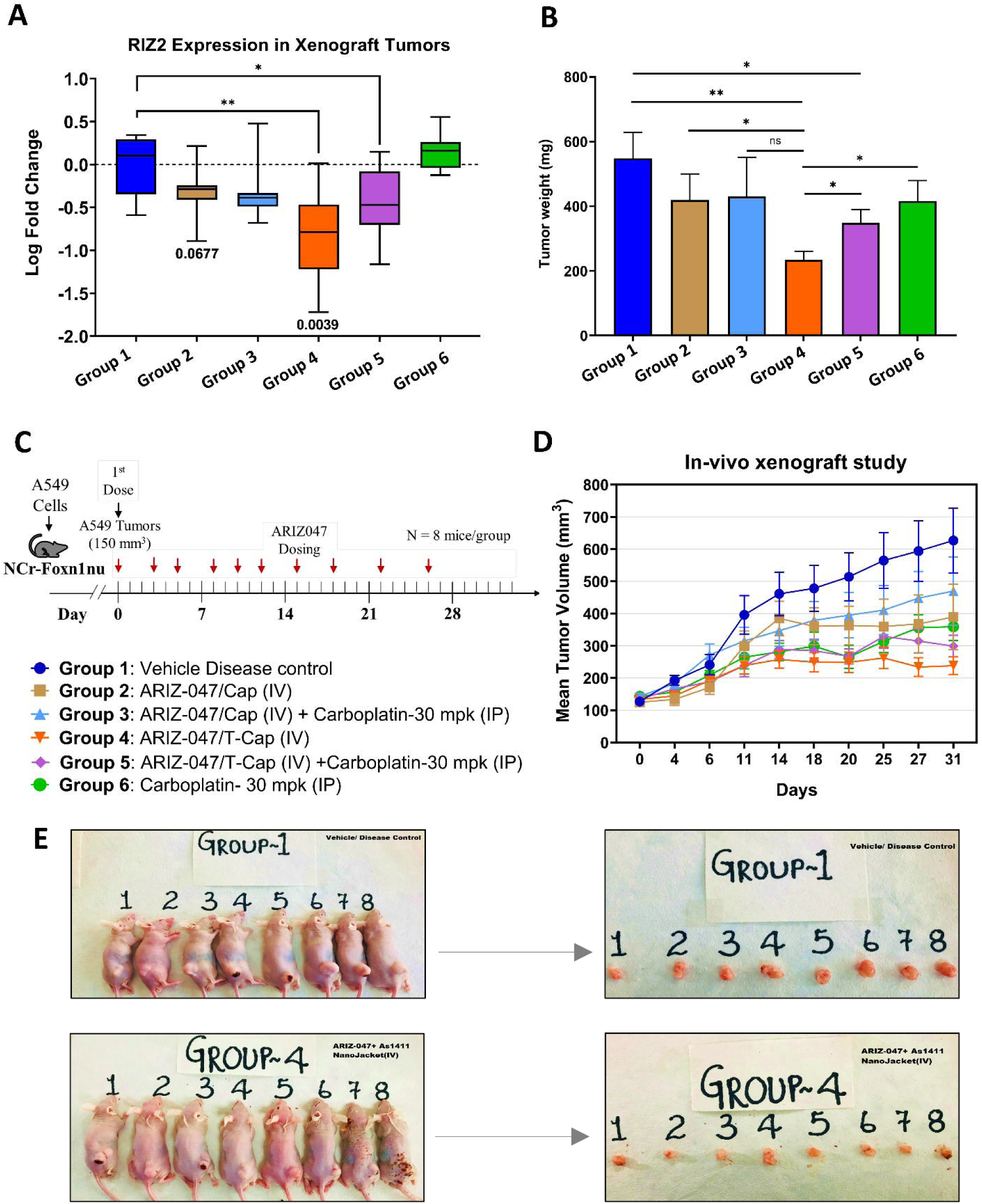
**A.** Comparison of RIZ2 expression level in Xenograft tumor with the treatment of vehicle control, ARIZ-047/Cap (IV), ARIZ-047/Cap (IV) + Carboplatin-30 mpk (IP), ARIZ-047/ T-Cap (IV), ARIZ-047/T-Cap (IV) + Carboplatin-30 mpk (IP), and Carboplatin-30 mpk (IP) (**p<0.005, *p<0.05). **B.** A549 tumor xenograft study reveals treatment with ARIZ-047 siRNA loaded nucleolin targeting ligand nanoparticles at 1mg/kg leads to an 80% decrease in mean tumor volume compared to vehicle control (**p<0.005, *p<0.05). **C.** Timeline showing the Schedule of ARIZ-047 dosing into the tumor-induced NCR-Fox-n1nu xenograft mice and description of different mice groups. **D.** Comparison of tumor weight with the treatment of vehicle control, ARIZ-047/Cap (IV), ARIZ-047/Cap (IV) + Carboplatin-30 mpk (IP), ARIZ-047/T-Cap (IV), ARIZ-047/T-Cap (IV) + Carboplatin-30 mpk (IP), and Carboplatin-30 mpk (IP). Tumor weights were compared using the Kruskal-Walli’s test (p = 0.0123) and Dunn’s multiple comparisons test. A statistically significant difference was observed between the vehicle control and ARIZ-047/T-CaP (IV) group (*p = 0.0140). No other pairwise comparisons showed significant differences (p > 0.05). **E.** Comparison of Xenograft tumor size from the group treated with vehicle control (group 1) and ARIZ-047/T-CaP (IV) (group 4).

## Discussion

In this study, we validated our in-house developed siRNA therapeutic molecule termed ARIZ-047, encapsulated with a nanojacket of calcium phosphosilicate, targeting an oncogenic RIZ2 gene. The initial objective of this study was to investigate the expression profiles of RIZ1 and RIZ2 in the lung cancer cell line compared to the normal cell line and understand their potential to be used as therapeutic targets. The transcript analysis using GEPIA2 and our experimental validation confirmed significant downregulation of RIZ1 and upregulation of RIZ2 in lung adenocarcinoma cells, suggesting their role in tumor progression. Several studies have also reported the downregulation of RIZ1 but not RIZ2 in various forms of cancers related to the breast, skin, colon, liver, and bone, including the lung (31–33). This discrepancy highlights that the RIZ variants might play distinct roles in tumorigenesis, where RIZ2 can be used as a potential target for inhibiting metastasis in lung cancer. Existing treatments, such as chemotherapy, radiotherapy, immunotherapies (CAR-T, and antibody therapy) (34–36), and targeted therapies for EGFR (37), ROS1 (38), RET (39), MET (40,41), BRAF (42), and KRAS (43,44) mutations have limitations such as toxicity, cost, and drug resistance. The targeted therapies developed against these mutated genes are effective for a small percentage of patients, and over the years, the cancer cells have developed resistance against the therapies available (45). However, the dysregulation of PRDM2 isoforms is disrupted in over 70% of patients, making ARIZ-047 potentially more effective than the available targeted therapies. The treatment with ARIZ-047 led to a change in the expression level of RIZ1 and RIZ2 without affecting the off-target genes ATP1A4 and TOPBP1. The specificity of ARIZ-047 was also supported by the cell viability experiments where ARIZ-047 significantly targeted cancer cell lines without harming normal cells. This highlights the potential of ARIZ-047 as a targeted therapeutic agent that could overcome the limitations and side effects associated with conventional chemotherapy and radiotherapy. The specificity of ARIZ-047 was also compared to a widely used chemotherapeutic drug, i.e., cisplatin, which interestingly did not show a significant reduction in A549 viability but significantly affected the viability of HBE1. The comparison of ARIZ-047’s efficacy with cisplatin suggests its potential superiority in selectively targeting cancer cells. Cisplatin interferes with the DNA replication and transcription machinery and imparts anti-tumor effects; however, cisplatin was also reported to lose efficacy due to the drug-resistance properties of tumor cells, leading to tumor progression (46,47). Transcriptome analysis revealed 111 DEGs upon ARIZ-047 treatment, significantly impacting the JAK-STAT and Wnt signaling pathways-both crucial for oncogenesis (48–50).

The elevated gene expression within the Wnt pathway during ARIZ-047 treatment could represent a compensatory mechanism by the tumor cells, which might be an attempt to regain their oncogenic potential. Conversely, the upregulated genes might also indicate a counterintuitive mechanism where the overexpression of certain genes within the Wnt signaling pathway could have suppressive effects on tumorigenesis.

A comprehensive protein-protein interaction (PPI) network analysis was executed in the presence and absence of RIZ2. Deletion of RIZ2 led to a notable shift in global connectivity patterns for the DEGs involved in numerous biological pathways. The PPI network analysis upon RIZ2 reintroduction demonstrated RIZ2’s critical interactions with key regulatory proteins such as PRDM1, ZFP36, DUSP1, and MCL1. PRDM1 regulates both B and T cell differentiation and plays a role in immunosuppression (51). By inhibiting RIZ2, ARIZ-047 may reduce immunosuppression and enhance anti-tumor immunity, particularly where RIZ2 amplifies PRDM1’s function. Additionally, PRDM1 downregulation has been linked to increased invasion and metastasis in lung cancer (52), suggesting RIZ2 may modulate PRDM1’s transcriptional activity. MCL1, an anti-apoptotic protein, contributes to tumor survival (53,54). RIZ2’s direct interaction with MCL1 suggests a role in regulating apoptosis and tumor progression, making its inhibition by ARIZ-047 a potential strategy to restore the apoptotic pathway. Similarly, ZFP36, which is also known for its apoptotic role (55), interacts directly with RIZ2, which may influence the fate of cancer cells. Notably, the loss of ZFP36 leads to upregulation of BARX1, promoting proliferation, migration, and invasion of NSCLC cells (56). By inhibiting RIZ2, ARIZ-047 may help restore ZFP36 function, thereby reducing BARX1 expression and restraining cancer progression. DUSP1 is known to be elevated in various cancers, including lung cancer (57), where it suppresses the JNK signaling pathway, a key apoptosis regulator. The direct interaction between RIZ2 and DUSP1 may promote tumor survival, inhibiting JNK-mediated apoptotic pathways. By targeting RIZ2, ARIZ-047 may disrupt this interaction, potentially restoring JNK signaling and enhancing apoptosis.

The 3D spheroids used in this study simulated the in vivo architecture and provided a robust platform to assess the intricate interplay between RIZ2 and the Wnt signaling pathway. The formation of A549 spheroids was hampered on ARIZ-047 treatment, which was restored upon co-treatment with a Wnt signaling modulator, i.e., R-spondin, indicating the inhibitory effect of treatment on Wnt signaling. Interestingly, the robust growth observed in spheroids overexpressing RIZ2-GFP declared its role in stimulating cellular proliferation. We have also shown previously that spheroids overexpressing RIZ2 protein increase significantly in size (16). However, treatment with the Wnt-agonists, particularly LGK-974, blunted this growth significantly, underlying the intricate balance between RIZ2 and Wnt signaling modulation.

*In vivo* xenograft studies confirmed ARIZ-047’s efficacy, significantly reducing tumor size and RIZ2 expression at lower doses than carboplatin. Carboplatin is a platinum-based anti-cancer drug that has played a vital role in cancer therapy with satisfactory efficacy but also possesses serious side effects that limit its usage (58). ARIZ-047 exhibited substantial efficacy at lower doses with potentially fewer side effects than carboplatin. The reduction in tumor size between untreated mice and those treated with ARIZ-047 further supports our quantitative findings. Additionally, the contrast in tumor volumes between these groups visually captures the profound anti-tumor effects of ARIZ-047. In conclusion, the evident therapeutic potential of ARIZ-047 in vitro and *in vivo* offers promising avenues for lung cancer treatment, potentially offering an alternative to existing therapies.

## Data availability

The data supporting this study’s findings are available on request from the corresponding author. The dataset analyzed during the current study is available in the NCBI repository under the BioProject ID PRJNA1209369 (https://www.ncbi.nlm.nih.gov/bioproject/PRJNA1209369).

## Ethical approval

Animal studies were conducted by Sphaera Pharma Pvt. Ltd. Sphaera’s CPCSEA registration number is 1285/PO/RcBi/S/09/CPCSEA. The protocol for the xenograft study was IAEC_2022/05.

## Authors’ Contributions

**S. Kumar:** Investigation, Conceptualization, Data generation, analysis, writing-original draft, Review and editing. **M. Z. Malik:** Investigation, Conceptualization, Data generation, analysis, writing-original draft, Review and editing. **M. Chaturvedi:** Data generation and analysis, Review. **M. Mishra:** Data generation and analysis. **M. D. Donato:** Investigation, writing-original draft. **A. Casamassimi:** Investigation, writing-original draft. **C. Abbondanza:** Conceptualization, writing-review and editing. **P. Gazzerro:** Supervision, resources, writing-review, and editing. **C. Nguyen:** Experimented, Data generation, and analysis. **N. N. Nuñez:** Experimented, Data generation, and analysis. **S. Sen:** Animal experiment and data interpretation. **B. Niles:** Conceptualization, resources, investigations, writing-original draft, writing-review and editing, supervision, Validation, Data generation, and analysis. **R. Chaturvedi:** Conceptualization, resources, investigations, writing-original draft, writing-review and editing, supervision, Validation, Data generation, and analysis

## Acknowledgments

The authors thank the Council of Scientific and Industrial Research (CSIR), Govt. of India, for providing a research fellowship (to S. Kumar) during the senior research fellow’s tenure. The authors would also like to acknowledge the Advanced Instrumentation Research Facility (AIRF), Jawaharlal Nehru University, New Delhi, India, for providing the computational resources. A graphical abstract was created using icons and templates retrieved from <u>BioRender.com</u> with appropriate publication licenses (Agreement number: ID27TYQ2R3). The authors would also like to acknowledge Dr. Erika Di Zazzo, PhD (University of Molise), Dr. Monica Rienzo, PhD (University of Campania), and Prof. Giovanni Nassa (University of Salerno), for their valuable contributions to the discussions of the data and Bioinformatics analysis of the regulated PRDM2 genes.

## Authors’ Discloures

**C. Abbondanza** reports a funding source from the University of Campania, “Luigi Vanvitelli” D.R. n. 797 of 01/08/2024 “RIZnF”. **M. D. Donato** reports a funding source from the Ministry of University -Italy-P.R.I.N.2022-PNRR (P2022BWAJN_002 to MDD). ARIZ Precision Medicine supported this research. No disclosures were reported by other authors.

## Supplementary Figures

**Supplementary Figure 1:**
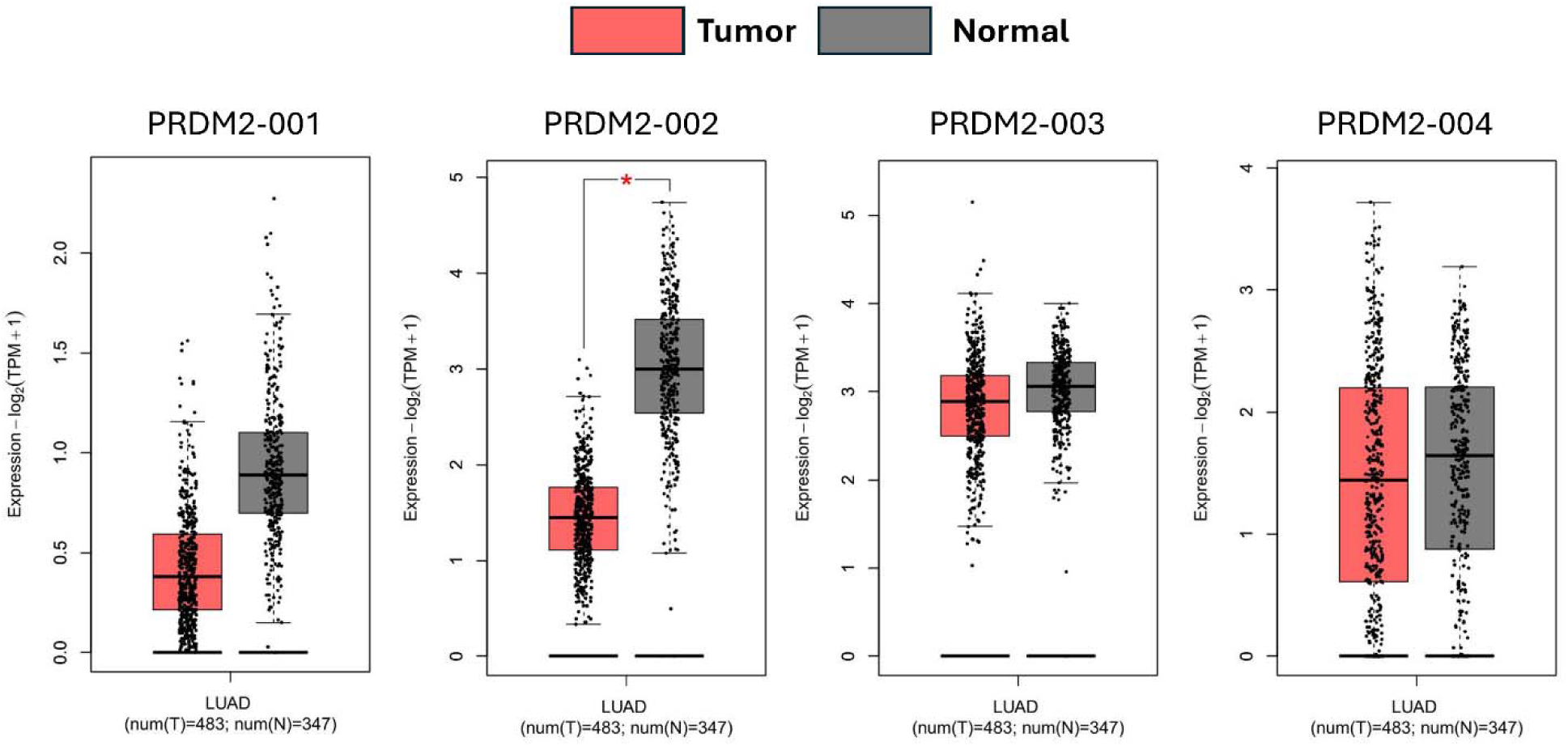
Expression profiles of PRDM2 transcripts in human tumor and normal tissues. The expression profiles of PRDM2 transcripts were analyzed using Gene Expression Profiling Interactive Analysis (GEPIA2). PRDM2-001 and PRDM2-002, representing the tumor suppressor variant RIZ1 transcribed from promoter 1, showed significant downregulation in tumor samples compared to normal tissues. In contrast, PRDM2-003 and PRDM2-004, corresponding to the oncogenic variant RIZ2 transcribed from promoter 2, exhibited consistent expression levels between tumor and normal samples. Data are presented as fold change in transcripts per million (TPM), normalized to β-actin.

**Supplementary Figure 2:**
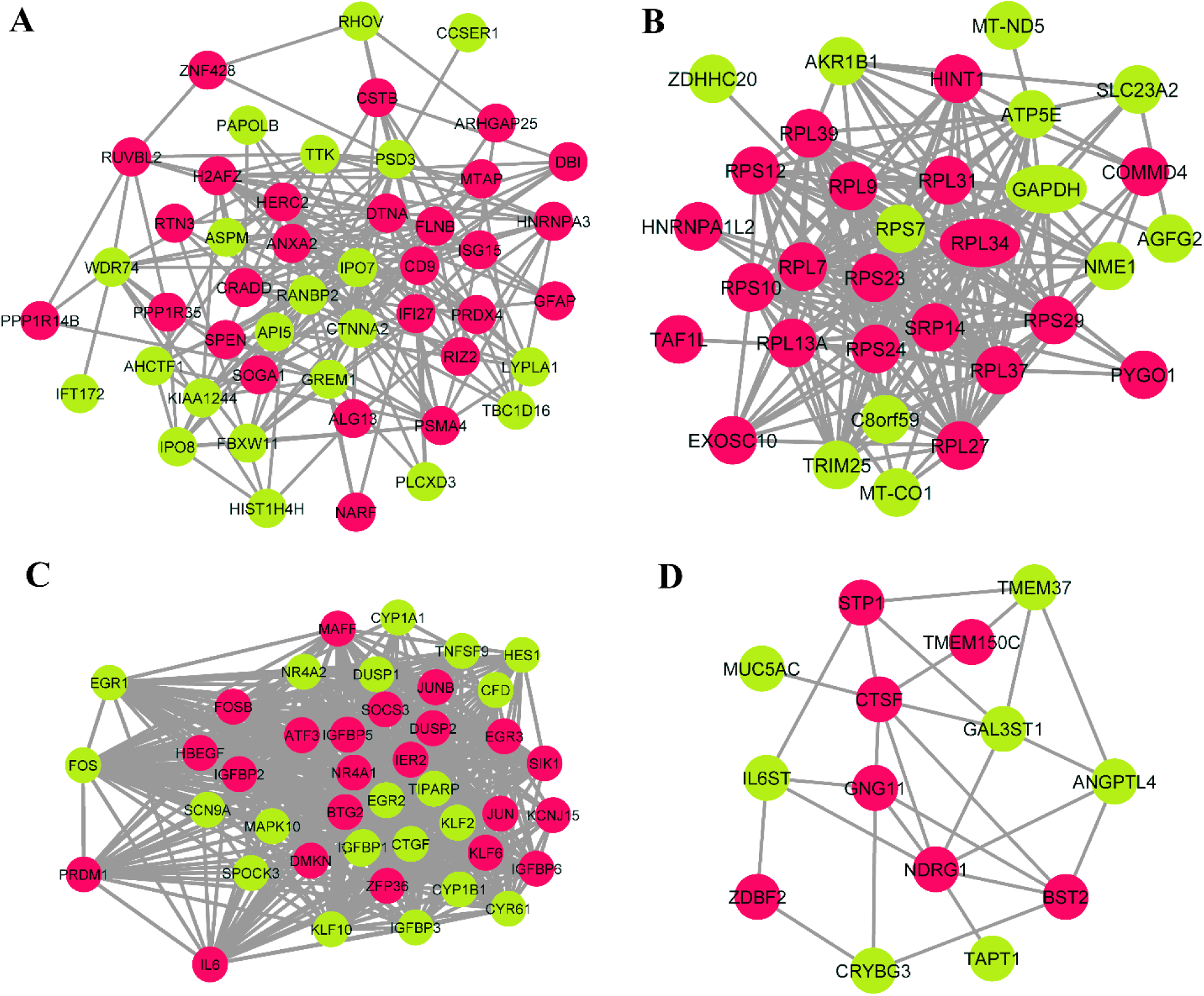
Protein-protein interaction (PPI) network of ARIZ047 responsive differentially expressed genes in A549 cells reveled by functional protein association network (STRING) analysis. **A.** Protein enriched in biological process involved in biological binding, **B.** Protein enriched in Biological process involved in Translation and Transcription, **C.** Protein enriched in Biological process involved in Cancer associated Signalling, **D.** Protein enriched in Biological process involved in Metabolic pathways and immune response.

**Supplementary Figure 3:**
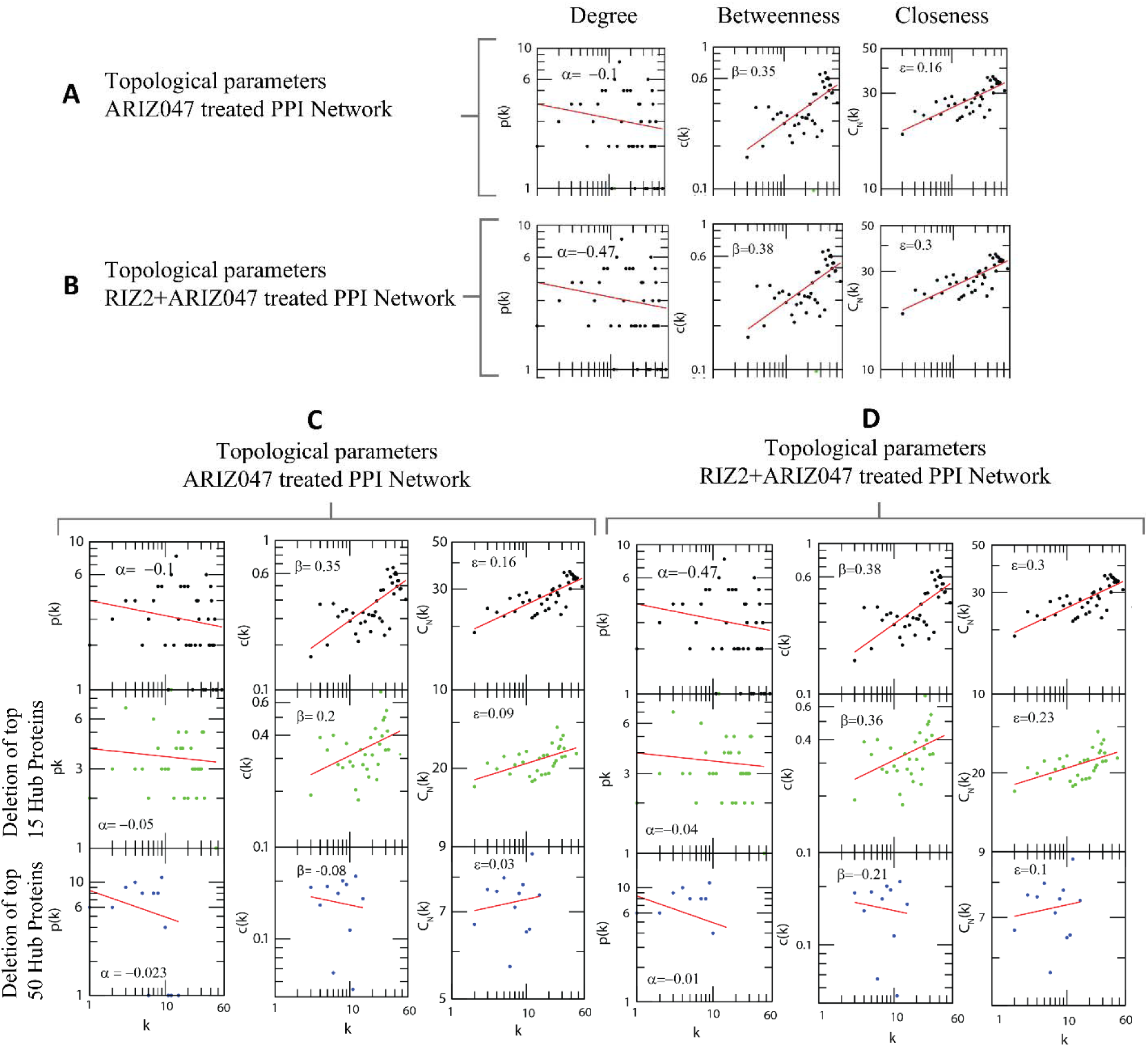
Topological parameters of protein-protein interaction network in response to treatment with: A. ARIZ047 treatment, B. ARIZ047 + RIZ treatment, C. Topological parameters of ARIZ047 treated protein-protein interaction network with deletion of top 15 and 50 hub proteins, D. Topological parameters of ARIZ047 + RIZ treated protein-protein interaction network with deletion of top 15 and 50 hub proteins.

